# DroNc-Seq: Deciphering cell types in human archived brain tissues by massively-parallel single nucleus RNA-seq

**DOI:** 10.1101/115196

**Authors:** Naomi Habib, Anindita Basu, Inbal Avraham-Davidi, Tyler Burks, Sourav R. Choudhury, François Aguet, Ellen Gelfand, Kristin Ardlie, David A Weitz, Orit Rozenblatt-Rosen, Feng Zhang, Aviv Regev

**Affiliations:** Broad Institute of MIT and Harvard, Cambridge MA 02142; McGovern Institute, Department of Brain and Cognitive Sciences, Massachusetts Institute of Technology, Cambridge MA 02140; John A. Paulson School of Engineering and Applied Sciences, Harvard University, Cambridge, MA 02138; Department of Physics, Harvard University, Cambridge, MA 02138; Howard Hughes Medical Institute, Department of Biology, Koch Institute of Integrative Cancer Research, Massachusetts Institute of Technology, Cambridge MA 02140

## Abstract

Single nucleus RNA-Seq (sNuc-Seq) profiles RNA from tissues that are preserved or cannot be dissociated, but does not provide the throughput required to analyse many cells from complex tissues. Here, we develop DroNc-Seq, massively parallel sNuc-Seq with droplet technology. We profile 29,543 nuclei from mouse and human archived brain samples to demonstrate sensitive, efficient and unbiased classification of cell types, paving the way for charting systematic cell atlases.

---

Single cell RNA-seq has become an instrumental approach to interrogate cell types, dynamic states and functional processes in complex tissues^1,2^. However, current protocols require the preparation of a single cell suspension from fresh tissue, a major roadblock in many cases, including clinical deployment, handling archived materials and application in tissues that cannot be readily dissociated. In particular, in the adult brain, harsh enzymatic dissociation harms the integrity of neurons and their RNA, biases data in favour of recovery of easily dissociated cell types, and can only be used on samples from young organisms, precluding, for example, those obtained from deceased patients with neurodegenerative disorders. To address this challenge, we^3^ and others^4^ developed single nucleus RNA-seq (*e.g.*, sNuc-Seq^3^ and Div-Seq^3^) for analysis of RNA in single nuclei from fresh, frozen or lightly fixed tissues. sNuc-Seq can handle even minute samples of complex tissues that cannot be successfully dissociated, and provide access to archived or banked samples, such as fresh-frozen or lightly fixed samples. However, it relies on sorting nuclei by FACS into plates (96 or 384 wells), and thus cannot easily be scaled to profiling tens of thousands of nuclei (such as human brain tissue) or large numbers of samples (such as tumor biopsies from a patient). Conversely, massively parallel single cell RNA-seq methods, such as Drop-Seq^5^, InDrop^6^ and related commercial tools^7,8^ can be readily applied at this scale^9^ in a cost-effective manner^10^, but require a single cell suspension as input.

To address this challenge, we developed DroNc-seq (**Fig. 1a**), a massively parallel single nucleus RNA-seq method that combines the advantages of sNuc-Seq with the scale of droplet microfluidics to profile thousands of nuclei at very low cost and massive throughput. DroNc-Seq was modified from Drop-Seq^5^ to accommodate for the smaller size and relatively lower amount of RNA in nuclei compared to cells. Specifically, we modified the microfluidics design (**Supplementary Fig. S1A, B**) to generate smaller coencapsulation droplets (75 μm diameter) and flow parameters; we optimized the nuclei isolation protocol to reduce processing time and increase capture efficiency (**Supplementary Fig. S1C**); and we changed the downstream PCR conditions (**Methods**). We validated for single nucleus specificity using species-mixing experiments^5^, in which we combine nuclei from human 293 cells and mouse 3T3 cells in one DroNC-seq run, to assess single nucleus purity, as previously performed for cells^5^ (**Supplementary Fig. S1D**). Notably, the DroNc-Seq device and workflow are compatible with current Drop-Seq platforms.

**Figure 1.**
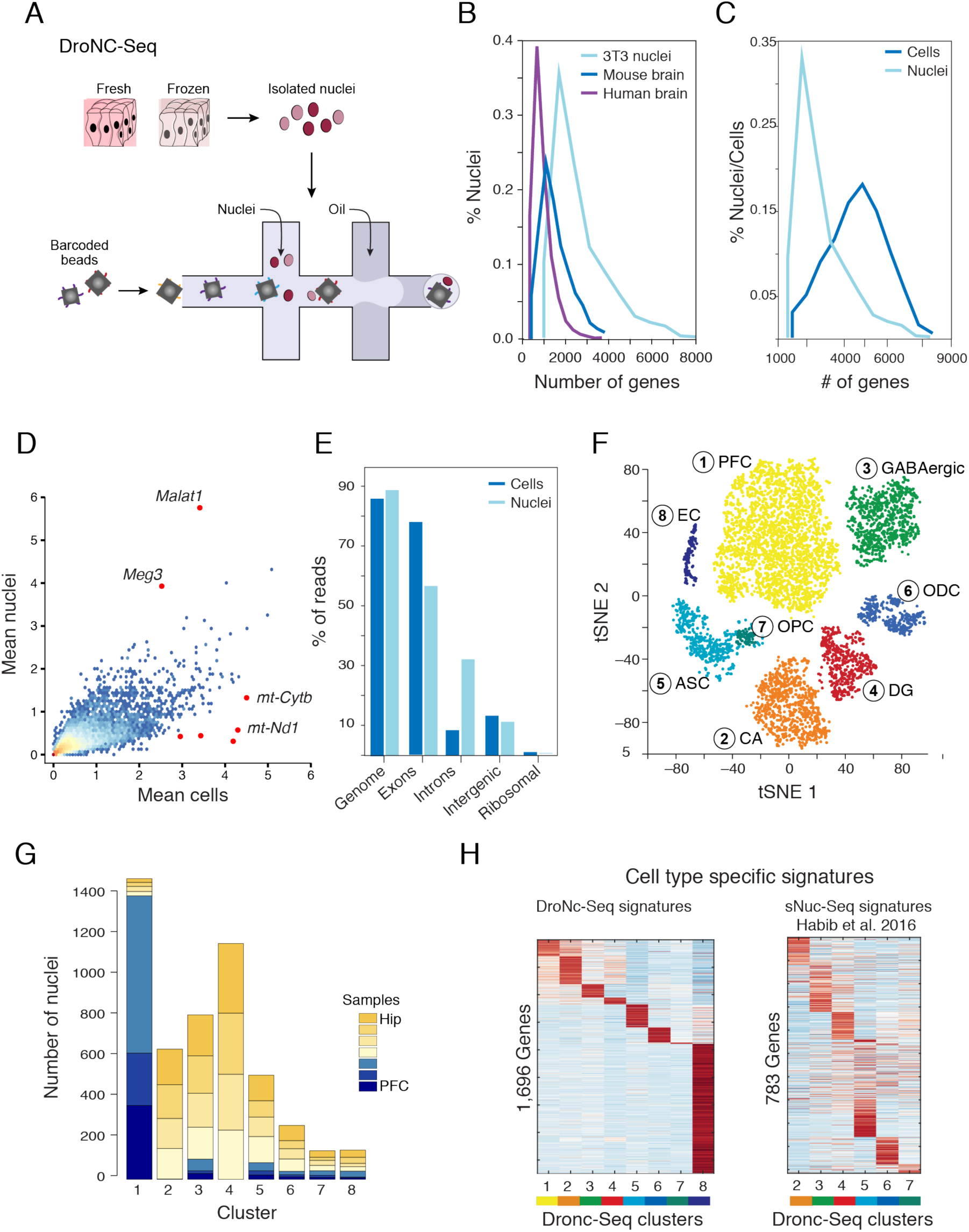
DroNc-Seq: Massively parallel single nucleus RNA-Seq. **(a)** Overview of DroNc-Seq. (**b-e**) Quality measures. (**b**) Distribution of number of genes detected (X axis) in DroNc-Seq of nuclei isolated from 3T3 mouse cells line, mouse frozen brain tissue, and human frozen archived brain tissue (**Methods**). **(c)** Distribution of number of genes detected per 3T3 cell (by Drop-Seq) or nucleus (by DroNc-Seq). (**d**) The percent of reads (Y axis) mapped to the: genome, exons, introns, intergenic regions and rRNA loci (X axis) of the mouse genome, for cells and nuclei. (**e**) Scatter plot comparing the average expression levels detected in single 3T3 nuclei (Y-axis, by DroNc-seq) and cells (X-axis, by Drop-Seq). Red dots mark outlier genes highly expressed in one but not the other experiment. (**f-h**) DroNc-Seq analysis of adult frozen mouse hippocampus (hip) and prefrontal cortex (PFC) brain regions. (**f**) A 2 dimensional t-stochastic neighbourhood embedding (tSNE) plot of 5,592 DroNc-Seq nuclei profiles from adult frozen mouse hippocampus (hip) (3 samples) and prefrontal cortex (PFC) (2 samples, each with >20,000 reads per nucleus), colored by clustering and labelled *post hoc* by cell types and anatomical distinctions (1. PFC = pyramidal neurons from the PFC, 2. CA = pyramidal neurons from the hip CA, 3. GABAergic = GABAergic neurons, 4. DG = granule neurons from the hip dentate gyrus (DG), 5. ASC = astrocytes, 6. ODC = oligodendrocytes, 7. OPC = oligodendrocyte precursor cells, 8. EC = endothelial cells). (**g**) Number of nuclei (Y axis) from each sample (PFC = blue gradient, hip = yellow gradient) associated with each cluster (X axis), showing that each cluster is supported by multiple samples, and most by both brain regions. (**h**) Signatures of differentially expressed genes. Right: The average expression in each DroNc-Seq cluster (column) of signature genes (**Methods**, rows) that are differentially expressed in the DroNc-Seq data for each cell type cluster derived from the DroNc-Seq data (numbered as is **f**). Expression is centred per row (color bar). Right: The average expression in each relevant DroNc-Seq cluster (numbered as is **f**, column) of signature genes previously identified on sNuc-Seq profiles^3^ for the corresponding cell types, showing that DroNc–seq captures similar diversity in nuclear RNA profiles between cell types.

DroNc-Seq robustly generated high quality expression profiles from nuclei isolated from a mouse cell line (3T3, 4,442 nuclei), adult mouse brain tissue (9,219), and adult human post-mortem frozen archived tissue (20,324 nuclei). It detected, on average 3,152 genes (6,614 transcripts) for 3T3 nuclei, 1,500 genes (2,614 transcripts) for nuclei from adult mouse brain, and 1,000 genes (1,337 transcripts) for nuclei from human post mortem brain tissue (**Methods**, **Fig. 1b**).

To assess Dronc-Seq’s throughput and sensitivity, we profiled the same 3T3 cell culture at both the single cell (with Drop-Seq) and single nucleus (with DroNc-Seq) levels, each sequenced to an average depth of 120,000 reads per nucleus or cell. Both methods yielded high quality libraries, detecting, on average, 4,770 and 3,152 genes for cells and nuclei, respectively (**Fig. 1c**). DroNc-Seq had somewhat reduced throughput, with 2,982 / 300,000 input nuclei passing filter (∼1%), compared to 5,175 / 100,000 cells (5%) passing filter per run. The average expression profile of single nuclei was well-correlated with the average profile of single cells (Pearson *r* = 0.87, **Fig. 1d**), albeit somewhat lower than the correlation between the average profiles of two replicates of Drop-Seq (*r* = 0.99) or DroNc-Seq (*r* = 0.99). Those genes with significantly higher expression in nuclei (*e.g*., the lncRNAs *Malat1* and *Meg3*) or cells (mitochondrial genes Mt-nd1, Mt-nd2, Mt-nd4, Mt-cytb) (**Fig. 1d**) were consistent with their known distinct enrichment in nuclear *vs*. non-nuclear compartments (**Supplementary Table 1**). Interestingly, while in both methods over 85% of reads align to coding loci, in cells 80% of these reads map to exons, whereas in nuclei 56% map to exons and 32% to introns (**Fig. 1e**), reflecting the enrichment of nascent, pre-processed transcripts in the nuclear compartment^3,11-14^.

Clustering^9^ of 5,592 nuclei profiled from frozen adult mouse hippocampus (3 samples) and prefrontal cortex (3 samples) (each with >20,000 reads per nucleus**, Methods**) revealed groups of nuclei corresponding to known cell types (*e.g.*, GABAergic neurons) and anatomical distinctions between the brain regions and within the hippocampus (*e.g.*, CA1, CA3, dentate gyrus; **Fig. 1f**). Neurons of the same class but from different brain regions (and different samples) group together, as was also the case for GABAergic neurons, glia and endothelial cells (**Fig. 1f-g**). Among the non-neural cells, different glia cell types, including astrocytes, oligodendrocytes and oligodendrocyte precursor cells (OPC), readily partitioned into separate clusters, despite their relatively low RNA levels and correspondingly lower numbers of detected genes (**Fig. 1f**). Finally, DroNc-Seq of mouse hippocampus compared well to sNuc-Seq of the same region^15^, maintaining the ability to detect the same cell types and correlated cell-types specific signatures (**Fig. 1h, Supplementary Table 2**) with increased throughput, despite a lower number of genes detected per nucleus in the massively parallel setting.

To demonstrate the utility of DroncSeq on archived human tissue, we profile adult (40-65 years old) human post-mortem frozen brain tissue archived by the GTEx project^16^. We analysed 10,368 nuclei (each with >20,000 reads per nucleus) from five frozen postmortem archived samples of adult human hippocampus and prefrontal cortex, revealing distinct nuclei clusters corresponding to the known cell types in these regions (**Fig. 2a**). We readily annotated each cell type cluster *post-hoc* by its unique expression of known canonical marker genes (**Fig. 2b**), including rare types, such as adult neuronal stem cells specifically found in the hippocampus (**Fig. 2a**, cluster 9). Although the human archived samples vary in the quality of the input material, DroNc-Seq yielded high-quality libraries of both neurons and glia cells from each sample (**Fig. 2c**, bottom), and each cluster was supported by multiple samples (**Fig. 2c**, top), demonstrating the robustness and utility of DroNc-Seq for clinical applications.

**Figure 2.**
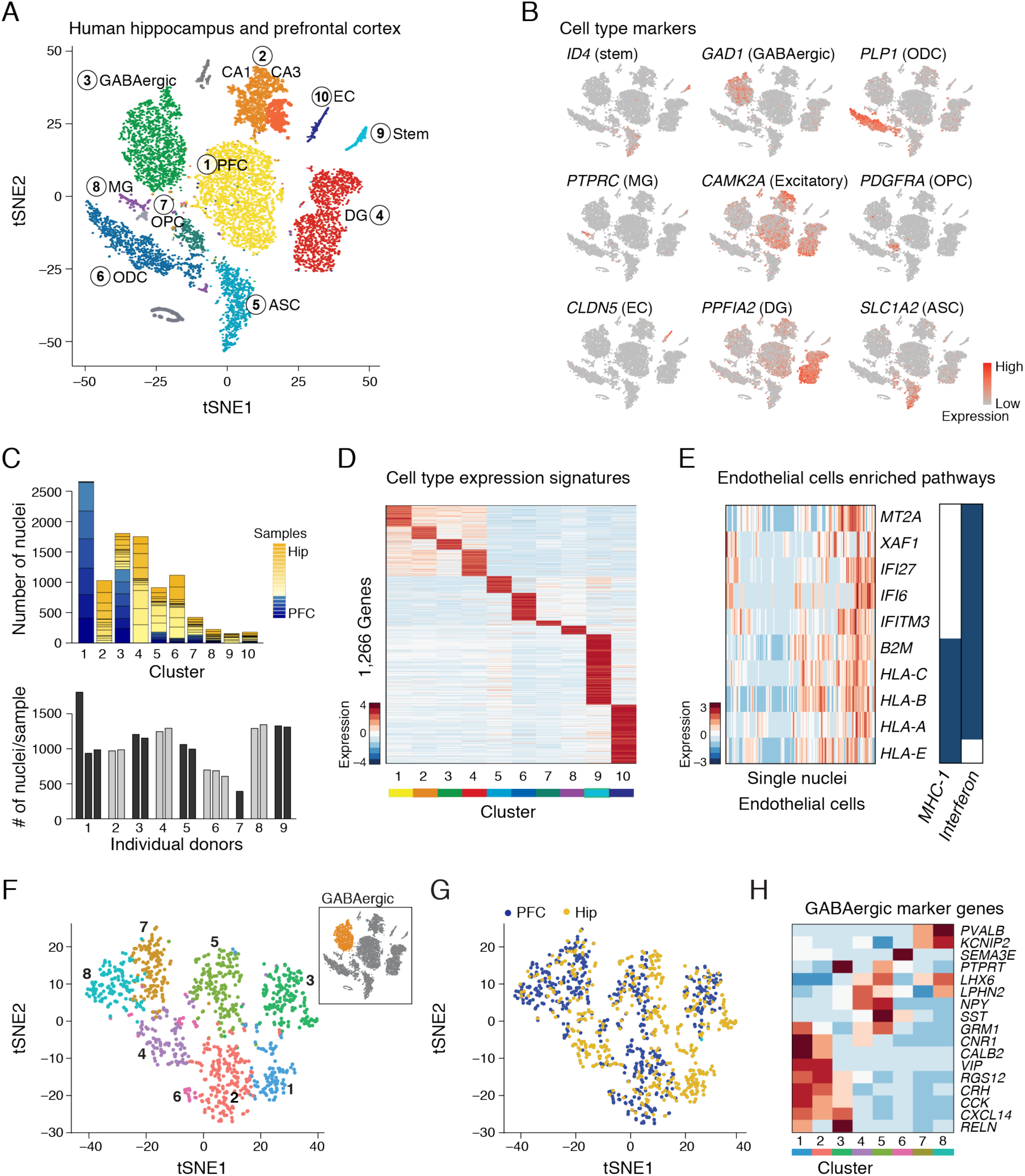
DroNc-Seq distinguished cell types and signatures in adult post-mortem human brain tissue. (**a**) Cell type clusters. tSNE embedding of 10,368 DroNc-Seq nuclei profiles from adult frozen human hippocampus and prefrontal cortex (PFC), each with >20,000 reads per nucleus. Clusters are color-coded and labelled *post-hoc* (1. PFC = pyramidal neurons from the PFC, 2. CA = pyramidal neurons from the hip CA, 3. GABAergic = GABAergic neurons, 4. DG = granule neurons from the hip dentate gyrus (DG), 5. ASC = astrocytes, 6. ODC = oligodendrocytes, 7. OPC = oligodendrocyte precursor cells, 8. MG = Microglia, 9. Stem = neuronal stem cells, 10. EC = endothelial cells) (**b**) Marker genes. Shown is the same plot as in (a) but with cells colored by the expression level of known cell type marker genes. (ID4-stem cells, GAD1 – GABAergic neurons, PLP1 – ODC, PTPRC – microglia, CAMK2 – excitatory neurons, PDGFRA – OPC, CLDN5 – EC, PPFIA2 – DG, SLC1A2 – ASC) (**c**) Successful DroNc-Seq across samples of different quality. Top: Number of nuclei (Y axis) from each sample (color code) associated with each cluster (X axis), showing that each cluster is supported by nuclei from multiple samples. Bottom: Number of nuclei passing quality filters (Y axis) recovered from each of 19 human tissue samples from 9 donors (X axis, sorted). (**d**) Cell type signatures. Heatmaps of the average expression of signature genes (rows; FDR <0.01) in nuclei in each of the clusters in (a). (**e**) Interferon signalling and MHC I genes in single endothelial cells. Shown is the expression of each gene across the nuclei in the endothelial cluster in (a). (**f-h**) Sub-types of GABAergic neurons. (**f,g**) tSNE embedding of DroNc-Seq nuclei profiles from the GABAergic neuronal cluster (in **a**), color coded by sub-clusters (f) or brain region (g). (**h**) Heatmap of the average expression of known marker genes of sub-types of GABAergic interneurons, in each of the nuclei sub-clusters in (f) (columns).

Finally, we determined cell-type specific gene signatures for each human cell type cluster (**Fig. 2d**), as well as a pan-neuronal signature, a pan-glia signature, and signatures for neuronal stem cells and endothelial cells (**Supplementary Table 3**). Signatures are enriched for key relevant pathways (FDR<0.01, **Methods)**. For example: Neuronal stem cells signatures are enriched for the expression of genes regulated by NF-kB in response to TNF signalling^17^; Endothelial cells are enriched for the expression of immune pathways, such as MHC I genes and interferon signalling (**Fig. 2e**), consistent with the known role of interferon signalling in modulation of the blood brain barrier^18^. Moreover, we captured finer distinctions between closely related cells (**Fig. 2f** and **Supplementary Fig. 2**), such as, distinct sub-types of GABAergic neurons (**Fig. 2f**), each robustly identified across biological replicates (**Supplementary Fig. 3a**), and often from both brain regions (**Fig. 2g**). Two of the GABAergic neuron sub-clusters are specific to the hippocampus (**Supplementary Fig. 3a, Fig. 2f**, clusters 1 and 4); these too are supported by multiple samples (**Supplementary Fig. 3a**). We associated each GABAergic neuron sub-cluster with a distinct combination of canonical markers (**Fig. 2h**), as previously reported in the mouse brain^3,19,20^.

In conclusion, DroNc-Seq is a massively-parallel single nucleus RNA-seq method, which is robust, cost-effective, and easy to use. Our results show that DroNc-Seq profiling from both mouse and human frozen archived brain tissues successfully identified cell types and sub-types, rare cells, expression signatures and activated pathways, opening the way to systematic single nucleus analysis of complex tissues that are either inherently challenging to dissociate or already archived. This will help create vital atlases of human tissues and clinical samples.

## Acknowledgements

We thank Karthik Shekhar, Christoph Muus and Eugene Drokhlyansky for helpful discussions, Timothy Tickle and Asma Bankapur for technical support, and Leslie Gaffney for help with graphics. Work was supported by the Klarman Cell Observatory, NIMH grant U01MH105960 and NCI grant 1R33CA202820-1 (to A.R.). Microfluidic devices were fabricated at the Center for Nanoscale Systems (CNS), Harvard University, member of the National Nanotechnology Coordinated Infrastructure Network (NNCI), and supported by the National Science Foundation under NSF award no. 1541959. A.R. is an Investigator of the Howard Hughes Medical Institute. A.R. is a member of the Scientific Advisory Board for Thermo Fisher Scientific, Syros Pharmaceuticals and Driver Genomics. F.Z. is supported by the NIH through NIMH (5DP1-MH100706 and 1R01-MH110049); NSF; the New York Stem Cell Foundation; the Howard Hughes Medical Institute; the Simons, Paul G. Allen Family, and Vallee Foundations; the Skoltech-MIT Next Generation Program; James and Patricia Poitras; Robert Metcalfe; and David Cheng. F.Z. is a New York Stem Cell Foundation-Robertson Investigator. D.A.W. thanks NSF DMR-1420570, NSF DMR-1310266 and NIH P01HL120839 grants for their support. NH is a Howard Hughes Medical Institute fellow for the Helen Hey Whitney Foundation. N.H., A.B., I.A.D., D.A.W., F.Z. and A.R. are inventors on international patent application PCT / US16 / 59239 filed by Broad Institute, Harvard and MIT, relating to method of this manuscript. GTEx is supported by the Common Fund of the Office of the Director of the United States National Institutes of Health, through Contract HHSN268201000029C (to K.A LDACC, Broad Institute).

## Supplementary Procedures and Methods

### Materials and Methods

#### Experimental procedures

##### Microfluidic device design and fabrication

Different microfluidic coflow devices were designed using AutoCAD (AutoDESK, USA) and tested using COMSOL Multiphysics as well as empirically. Microfluidic devices were fabricated using soft lithographic techniques^21^. Briefly, a negative mold of poly (dimethylsiloxane) elastomer (Sylgard 184, Krayden, Inc., Cat # DC4019862) was cast from a positive master mold made of SU8 photoresist (MicroChem, Cat # SU8-3050) via replica molding. The PDMS mold was then bonded to a 50 mm x 75 mm glass slide (Corning, Cat # 2947-75X50) using O_2_ plasma bonding, followed by baking the PDMS-glass device in an oven at 65°C for 1 hour. The device was then treated with Aquapel Glass Treatment (Aquapel, # 47100) to render all microfluidic channels hydrophobic to facilitate aqueous droplet generation in a continuous oil phase. The devices were extensively tested on a microfluidic setup consisting of an optical microscope and three syringe pumps (KD Scientific, Cat # KDS 910), using bare beads (Tosoh, Japan, Cat # HW-65s) in Drop-Seq Lysis Buffer (DLB^5^; a 10 ml stock consists of 4 ml of nuclease-free H_2_O, 3 ml 20 % Ficoll PM**-**400 (Sigma, Cat # F5415-50ML), 100 μl 20 % Sarkosyl (Teknova, Inc., Cat # S3377), 400 μl 0.5 M EDTA (Life Technologies), 2 ml 1M Tris pH 7.5 (Sigma), and 500 μl 1M DTT (Teknova, Inc., Cat # D9750), where the DTT is added fresh before every experiment) and 1x PBS, to optimize flow and bead occupancy parameters in drops. Droplet generation under different flow conditions was assessed in real time under an optical microscope (Olympus, Model # IX83) at 4x magnification using a fast camera (Photron, Model # SA5), and later by sampling the emulsion generated using disposable Neubauer Improved Hemocytometer (Life Technologies, Cat # 22-600-100) to check droplet integrity and size, as well as bead occupancy in drops. The device design is provided as a **Supplementary File 1**.

##### Cell culture

3T3 cells were cultured and prepared as described^5^. For DroNC-seq run cells were washed once with PBS, scrapped with 2ml Nuclei EZ lysis buffer (Sigma Cat # EZ PREP NUC-101) and processed as described below for tissues.

##### Dissection of and mouse hippocampus and cortex

Microdissections of the mouse hippocampus and prefrontal cortex regions were performed under a stereomicroscope as previously described^3^. Dissected sub-regions were flash frozen on dry ice and stored at-80°C until processed for nuclei isolation.

###### Human hippocampus and prefrontal cortex samples

Human hippocampus and pre-frontal cortex samples were collected as part of the Genotype-Tissue Expression (GTEx) project. All samples were collected from recently deceased postmortem, non-diseased donors as previously described^16,22^. Briefly, brains were collected for only a subset of donors when consent and conditions (donors could not have been on a ventilator for the 24 hours prior to death) allowed. All collected brains were immediately removed from the body and placed on wet ice, then shipped to the Brain Bank at the University of Miami. Up to 11 regions of brain were then carefully dissected by the brain bank upon receipt. All brain tissues sampled were placed in to cryovials and flash frozen in Liquid N2, then shipped to the LDACC at the Broad institute for processing and analysis. For this study, samples of frozen hippocampus and prefrontal cortex were selected from 5 male donors, ranging in age from 40-65. We used the quality of RNA derived from the tissues as a proxy for tissue quality, and selected tissues with corresponding RNA RIN values of 6.9 or higher (average RIN was 7.3). Average post-mortem ischemic interval of the brains collected was 12.4 hours.

##### Nuclei isolation

Nuclei were isolated by using the EZ nuclei isolation kit (Sigma, #EZ PREP NUC-101). Briefly, tissue samples were cut into pieces smaller than 0.5 cm, dounce homogenized in 2 ml of ice-cold Nuclei EZ lysis buffer and incubated on ice, for 5 minutes, with additional 2 ml of ice-cold Nuclei EZ lysis buffer. Nuclei were collected by centrifugation at 500 x g for 5 minutes at 4 °C, washed with 4 ml of ice-cold Nuclei EZ lysis buffer and incubated on ice for 5 minutes. After centrifugation, the nuclei preps were washed again in 4 ml of Nuclei Suspension Buffer (NSB; consisting of 1x PBS, 0.01% BSA and 0.1% RNAse inhibitor (Clontech, Cat #2313A)). Isolated nuclei were resuspended in 2ml of NSB, filtered through a 35 μm cell strainer (Corning, Cat # 352235) and counted. A final concentration of 300,000 nuclei/ml was used for DroNc-seq experiments.

##### Co-encapsulation of nuclei and barcode beads

A 10 μl sample of a single nuclei suspension prepared on ice in NSB (as described above) was stained with DAPI (ThermoFisher Scientific, Cat # D1306), loaded on a disposable Neubauer Improved Hemocytometer, and checked under the microscope to ensure that the nuclei are adequately isolated into singletons and no big clusters remain. The nuclei were also counted and suspended in NSB at a concentration of ∼300,000 nuclei/ml.

Barcoded beads (Chemgenes, Cat # Macosko-2011-10) were washed and filtered using 100 μm cell strainer (VWR, Cat #08-771-19), as previously described^5^. Because the DroNc-Seq microfluidic device has narrower channels (∼70 μm), they are more likely to clog in the presence of large beads. We therefore size-selected for beads that are less than 40 μm in diameter, using a 40 μm cell strainer (PluriSelect, Cat # 43-50040-03); these smaller beads consist of roughly 55% of the original population of barcoded beads. The barcoded beads were then suspended in Drop-Seq Lysis Buffer (DLB; described above) and counted using a disposable Fuchs-Rosenthal hemocytometer (VWR, Cat # 22-600-102) as previously described^5^, at concentrations ranging between 325,000 and 350,000 per ml.

The nuclei and barcoded bead suspension were then loaded into 3 ml syringes (BD Scientific, Cat # BD309695) connected to the custom-built DroNc-Seq microfluidic chip via 26G1/2 sterile needles (BD Scientific, Cat # BD305111) and PE2 tubing (Scientific Commodities, Inc. Cat # BB31695-PE/2), and flown at 1.5 ml/hr each, along with carrier oil (BioRad Sciences, Cat # 186-4006) at 16 ml/hr. This allowed us to co-encapsulate single nuclei and beads in ∼75 μm drops (vol. ∼ 200 pl) at 4,500 drops/sec and double Poisson loading concentrations. The smaller droplet volume in DroNc-Seq (at 75 μm diameter) results in a higher concentration of mRNA in these drops (more than 5x) compared to those used in Drop-Seq (at 125 μm diameter).

The theoretical Poisson loading concentration for devices with channels that are 75 μm deep and nominally 70 μm wide at 1/10 bead and nuclei occupancy is ∼520,000/ml, and that of 75 μm channel and 100 μm depth (also tested) is 340,000/ml. We tested bead and cell loading at this and other concentrations using species-mixing experiments^5^ (e.g., **Supplementary Fig. 1d**) and ease of bead flow as metrics and found that a bead concentration of 350,000/ml and a nuclei concentration of 300,000/ml had the best performance, in terms of low human-mouse doublet rate and fewer clogging events during droplet generation.

The barcoded beads are constantly stirred while loaded on the syringe pump, using a flea magnet (VP Scientific, cat # 782N-6-150) in the syringe and magnetic stirrer setup (VP scientific, Cat # 782N-3-150), also previously described^5^.

##### Droplet breakage, washes and RT

The resulting emulsion was collected via PE2 tubing into a 50 ml Falcon tube for a period of ∼22 min each, and left to incubate at room temperature for up to 45 min before proceeding to break droplets.

Emulsion collected after co-encapsulation had the droplets cream to the top with clear oil collected under the droplets. We carefully removed the excess clear oil, added 30 ml of 6x SSC (Teknova, Inc., Cat #, S0282) into each 50 ml Falcon collection tube, agitated it vigorously, and added 1 ml of 1H,1H,2H,2H-Perfluorooctan-1-ol (SynQuest Laboratories, Cat # 647-42-7). The tubes were again vigorously shaken by hand and centrifuged at 1000xg for 1 min. The supernatant was then carefully removed from each tube and an additional 30 ml of 6x SSC was added vigorously to kick up the beads from the oil-water interface into the aqueous phase. The beads that were kicked momentarily into the SSC were quickly removed with a 25 ml pipette and transferred into a clean 50 ml Falcon tube, leaving the heavier oil behind. The newly transferred beads and SSC mix were centrifuged again at 1000xg for 1 min. The supernatant was carefully removed leaving ∼ 1ml of SSC and bead sediment behind. This remaining SSC and bead mix was then carefully transferred into a 1.5 ml micro-centrifuge tube (Ambion, Cat # AM12450) and re-spun on a desktop micro-centrifuge for ∼ 20 sec that generated a substantial bead pellet. We removed any residual oil that got transferred into the 1.5 ml tube with a p200 pipette with low-retention pipette tip, although we found that any such residual oil did not appreciably affect Reverse Transcription (RT) efficiency. The beads in each tube were washed again in 1.5 ml of 6x SSC followed by another wash in 200 μl of 1x Maxima H-RT buffer (Fisher, Cat # EP0753). It is recommended to perform all washes and temporary storage of beads on ice. A pellet of barcoded beads in each microcentrifuge tube should have ∼ 130,000 beads.

We made a fresh batch of 200 μl RT mix for each barcoded bead aliquot, consisting of: 80 μl H2O, 40 μl Maxima 5x RT Buffer, 40 μl 20% Ficoll PM-400 (Sigma, Cat # F5415-50ML), 20 μl 10 mM dNTPs (Takara Bio, Cat # 639125), 5 μl RNase Inhibitor (Lucigen, Cat # 30281-2), 10 μl Maxima H-RT enzyme (Fisher, Cat # EP0753), and 5 μl 100 μM Template Switch Oligo, AAGCAGTGGTATCAACGCAGAGTGAATrGrGrG (IDT, custom RNA oligo). After all supernatant was carefully removed from each bead pellet, 200 μl of the above RT mix was added into each tube, and incubated under gentle rocking or tumbling, for 30 min at room temperature and then at 42 °C for 1.5 hr in a rotisserie-style hybridization oven for a total of two hours.

##### Post RT wash, exonuclease I treatment and PCR

Post RT, each barcoded bead has cDNA barcoded with the bead’s unique barcode (or BC) bound onto it, also referred to as a STAMP^5^. Each STAMP pellet was washed with (**1**) TE buffer containing 0.5% SDS (TE-SDS), once; (**2**) TE buffer containing 0.01% Tween-20 (TE-TW), twice; and (**3**) 10 mM Tris pH 8.0, once. The STAMPs were then treated with exonuclease I (New England Biolabs, Cat # M0293L) to remove all unextended primers. This was followed with another round of the above mentioned TESDS, and TE-TW washes, followed by a round of wash in DI water. Beads from multiple collections of a given sample were pooled at this point, resuspended in 1 mL of H_2_0, and counted, by mixing 10 μl of bead suspension with an equal volume of 20% PEG solution. Aliquots of 5,000 beads, resuspended in a PCR mix each consisting of 24.6 μl H_2_O, 0.4 μl 100 μM SMART PCR primer, AAGCAGTGGTATCAACGCAGAGT (IDT, custom DNA oligo), and 25 μl 2x Kapa HiFi Hotstart Readymix (Kapa Biosystems, Cat # KK2602), were amplified in separate wells on a skirted PCR plate, using the Eppendorf Thermocycler (Part # EP-950030020), with the following PCR steps: 95 °C for 3 min; then four cycles of: 98 °C for 20 sec, 65 °C for 45 sec, 72 °C for 3 min; then 10 cycles of: 98 °C for 20 sec, 67 °C for 20 sec, 72 °C for 3 min; and finally, 72 °C for 5 min. Amplified PCR product were pooled in batches of 16 wells (for mouse samples) or 32 wells (for human samples), each well consisting of the 5,000 STAMP aliquots, combined in a 1.5 ml Eppendorf tube, and cleaned with 0.6X SPRI beads (Ampure XP beads, Beckman Coulter, Cat # A63881). Whole transcriptome amplified (WTA) and Nextera libraries from mouse cortex, human cortex and human hippocampus generated libraries of different sizes, summarized in **Supplementary Table 1**.

| Sample | # WTA<br>PCR<br>wells<br>pooled | WTA<br>avg<br>peak<br>[bp] | WTA<br>min/max<br>[bp] | NTA<br>avg<br>peak<br>[bp] | NTA<br>min/max<br>[bp] | Illumina<br>lib. conc.<br>[pM] |
| --- | --- | --- | --- | --- | --- | --- |
| Mouse | 16 | 2148 | 617.33 /<br>8070 | 462 | 189 /<br>1557.4 | 2.2 |
| Human Cortex | 32 | 874 | 436 / 2520 | 435 | 185 / 1366 | 2.3 |
| Human<br>Hippocampus | 32 | 838 | 418.5/2408 | 437 | 244 / 846 | 2.3 |

##### WTA library QC and Nextera library prep

Purified cDNA was quantified using a Qubit dsDNA HS Assay kit (ThermoFisher Scientific, Cat # Q32854) and a BioAnalyzer High Sensitivity Chip (Agilent, Cat # 5067-4626). 550 pg of each sample library was fragmented, tagged and amplified using the Nextera XT sample prep kit (Illumina), and custom primer that enable selective amplification of the 3’ end (AATGATACGGCGACCACCGAGATCTACACGCCTGTCCGCGGAAGCAGTGGT ATCAACGCAGAGT*A*C, (IDT, custom DNA oligo)), according to the manufacturer’s instructions. The Nextera libraries are quantified again with Qubit dsDNA HS Assay kit and BioAnalyzer High Sensitivity Chip (See **Supplementary Table 1** for Nextera library sizes for human and mouse samples).

##### Sequencing

The libraries were sequenced at 2.2 pM (mouse) and 2.3 pM (human) on an Illumina Next-Seq 500 sequencer. We use Next-Seq 75 cycle v3 kits to sequence 20 bp and 65 bp paired end reads, with Custom Read1 primer, GCCTGTCCGCGGAAGCAGTGGTATCAACGCAGAGTAC (IDT, custom DNA oligo), according to Illumina loading instructions. The sequencing cluster density and percent passing filter number from different experiments vary according to the quality of nuclei samples used, but were optimized at around a cluster density of 220 and a 90% passing filter.

#### Computational data analysis

##### Preprocessing of DroNc-Seq data

###### Read filtering and alignment

Paired-end sequence reads were processed mostly as described before^5,9^. Briefly, the left read was used to infer the cell of origin based on the first 12 bases (the nucleus barcode or NB), and the molecule of origin based on the next 8 bases (Unique molecular Index or UMI). Reads were first filtered to remove all pairs where either the NB or the UMI had one or more bases with quality score less than 10. The right mate of each read pair (60 bp) was trimmed to remove any portion of the SMART adaptor sequence or large stretches of polyA tails (6 consecutive bp or greater). The trimmed reads were then aligned to the genome (mouse mm10 UCSC, human hg19 UCSC) using STAR v2.4.0a^23^ with the default parameter settings. Reads mapping to exonic regions of genes as per the mouse UCSC genome (version mm10) or the human UCSC genome (version hg19) were recorded. Exonic reads that mapped to multiple locations or to the antisense strand were discarded. Cell barcodes were checked for synthesis errors and corrected when needed (as previously described^9^).

###### Digital gene expression

Nucleus barcodes that represent genuine nuclei or cell RNA libraries from technical and sequencing noise were distinguished as previously described^5,9^ as true or “core” cell barcodes, by first ordering the nucleus barcodes by the total number of transcripts per nucleus barcode and estimating a cutoff based on a “shoulder” in the corresponding plot. Only nucleus barcodes larger than this cutoff were used in downstream analysis. We then collapsed nucleus barcodes by defining among these selected barcodes “core barcodes” or barcodes that other non-core barcodes can be collapsed with based on similarity (requiring at least 3,000 reads mapped to a barcode to qualify as a core barcode and using an edit distance of 1 to define similar barcodes). To account for amplification bias, we collapsed gene counts within each sample using their UMI sequences. The UMIs corresponding to all uniquely mapped sense reads (for a given gene) were recorded, and UMIs within an edit distance of 1 (substitutions only) were collapsed, as previously described^5^. The expression count for that particular gene in that particular nucleus was determined by counting the number of remaining unique UMIs. The data is assembled as a digital expression matrix (DGE) with genes as rows and nuclei as columns that served as the starting point for downstream analysis.

##### Gene detections and quality controls

###### Additional filtering of the expression matrix

The starting pool of nuclei was first filtered to remove nuclei based on their overall number of genes (minimum 100 genes). After this filtering we retained: (**1**) 1,710 cells from the 3T3 cell libraries (collected by Drop-Seq) across two replicates. (**2**) 4,442 3T3 nuclei libraries across 3 replicates. (3) 9,219 nuclei from the mouse brain (3,287 PFC nuclei from 3 mice and 5,932 hippocampus nuclei from 3 mice). (4) 20,324 nuclei from the human brain (8,057 PFC nuclei from 3 replicates and 12,267 hippocampus nuclei from 4 replicates). Note that for the species-mixing experiment we did not preform the additional filtering step, and we have 1,071 nuclei from two technical replicates. In downstream analysis we further tested gene detection thresholds for clustering robustness and applied thresholds on the number of reads associated with each nucleus barcode (20,000 reads for clustering analysis reported in the main figures). A gene is considered detected in a cell if it has at least two unique UMIs associated with it. For each analysis, we remove genes that were detected in less than 10 nuclei.

###### QC metrics

A list of quality metrics was obtained for the DroNc-seq single-cell libraries using Samtools (http://samtools.sourceforge.net/), Picard Tools (http://broadinstitute.github.io/picard/) and in-house scripts. For each single-nucleus library (identified based on its batch, replicate and a 12bp barcode), we calculated the total number of mapped reads (coding and UTR), the number of genes detected per cell, percentage of the total number of reads assigned to the nucleus barcode that were from: (**1**) coding regions, (**2**) UTRs, (**3**) intronic regions, (**4**) intergenic regions, (**5**) ribosomal RNA (rRNA), and (**6**) transcripts derived from the mitochondrial genome.

##### Comparison of Drop-Seq (cells) and DroNc-Seq (nuclei)

We compared between DroNc-Seq (nuclei) and Drop-Seq (cells) by several measures. (**1**) We calculated the capture rate, defined as the number of cells or nuclei recovered from a single run divided by the number of cells or nuclei loaded as input (300,000 for nuclei and 100,000 for cells): 1% for DroNc-seq and 5% capture for Drop-seq. (**2**) We compared the average and the distribution of the number of genes and transcripts detected for all cells and nuclei that pass our quality filter (**Figure 1**). (**3**) We compared the gene signatures of nuclei and cells, by computing the average expression signatures for each gene (mean of log UMI counts) in each replicate. We then computed the Pearson correlations between technical replicates of cells or nuclei (all have *r* = 0.99 +/stdev = 0.0023), and between nuclei and cells (*r* = 0.81 + /-stdev = 0.0024). (**4**) We tested for genes differentially expressed between cells and nuclei by pooling the technical replicates and testing for differential expression using student’s t-test, with FDR < 0.001, log-ratio > 1, and average expression across all nuclei or cell samples log(UMI count) > 3, which resulted in 2 genes up-regulated in the nuclei (lincRNAs Malat1 and Meg3), and 57 genes up regulated in cells, including many mitochondrial RNAs and ribosomal protein RNAs (known to be stable and thus enriched in cells compared to nuclei^11,12^), out of which the most significant ones are mt-Cytb and mt-Nd1; **Supplementary Table 1**). (**5**) We compared the fraction of the total number of reads that were mapped to (**1**) coding regions, (**2**) UTRs, (**3**) intronic regions, (**4**) intergenic regions, and (**5**) ribosomal RNA (as described above).

##### PCA, clustering and tSNE visualization

###### Dimensionality reduction using PCA

The dimension of the DGE matrix was reduced using principal component analysis (PCA), to project the original data to reduced linear dimensions where the most significant variance of the data is preserved, as determined based on the largest eigenvalue gap. We used the rpca function in R (package rsvd), which computes the top 100 principal components (or PCs) for efficiency. We then chose the most significant principal components (or PCs) to use as input in downstream analysis of clustering cells and embedding them into a 2-D space for visualization. For each separate run we chose the number of top PCs based on the largest eigen value gap. We found that the first PC is highly correlated with the number of genes detected in each nucleus (also when clustering individual cell type), as previously reported in many single cell analyses (Reviewed in Ref. ^1^), and is associated with highly and widely expressed genes. We thus excluded the first PC from the downstream analysis, which reduced the effects of the library quality and complexity on cluster identity.

###### Graph clustering

We partitioned the nuclei profiles into transcriptionally similar clusters using the top significant PCs as an input to a graph based clustering algorithm, as previously described^9^. In the first step, we compute a *k*-nearest neighbor (*k*-NN) graph on the data, where every nucleus is connected to each of its *k* nearest neighbors determined based on Euclidean distance in PC-space (using the nng function of the igraph package in R). We next used the k-NN graph as an input to the Infomap algorithm^24^, which decomposes an input graph into modules by deriving a compressive description of random walks on the graph. We ran Infomap using the cluster_infomap function in R. We compared the infomap clusters to an alternative clustering method, Louvain-Jaccard (using the cluster_louvain function in R) that receives as input the same *k*-NN graph as the infomap algorithm. Louvain clustering yielded a smaller set of clusters that highly overlap with the infomap clusters. The clustering results were visualized by coloring a tSNE 2-D map *post hoc* (below), but the tSNE embedding was not used to inform the clustering. We found that for detecting the major cell types, all clustering methods were robust and aligned with the visual separation seen in the tSNE embedding. We used the infomap algorithm and chose the *k* in the nearest neighbor graph that captured all the distinct point clouds in tSNE space using density clustering^25^ (across multiple tSNE runs for robustness), using k = 100 for clustering of over 1,000 nuclei and k = 50 otherwise

###### sub-clustering

To identify sub-types of cells we ran the same methods as described above on a specific subset of cells (one of the major clusters) to partition it to sub-clusters. For the human GABAergic sub-clusters, after the initial clustering step which separated the nuclei to distinct yet not well separated clouds of nuclei, we used the biSNE^3^ bi-clustering algorithm to choose genes that are localized in expression patterns in the tSNE mapping and thus are likely to be informative genes that are co-expressed by a close sub-set or sub-type of cells. We used these genes as an input to the same methods above (PCA, infomap clustering and tSNE visualization) to find finer distinctions between cells.

###### tSNE visualization

To visualize the nuclei, clustering and gene expression, we generated a two-dimensional (2D) non-linear embedding of the cells using t-distributed Stochastic Neighbor Embedding^26^ (tSNE). The scores along the top significant PCs estimated above were used as input to the algorithm. We ran the R implementation of tSNE, using the Rtsne package, with maximum of 1,000 iterations, disabling the initial PCA step and setting the perplexity parameter to 30 for detection of the major clusters and 25 for sub-clusters. Since tSNE can produce different visualizations in different runs, we used these coordinates for visualization (*i.e.*, not to identify cell clusters). The cells in the analysis of both the mouse and the human brain separated into distinct point clouds in tSNE space. This is true for any sub-set of cells that we chose from the data indicating the robustness of the clustering analysis. As we lower the threshold on the filtering step of nuclei from the DGE matrix we find that cells still partition to distinct clouds but the borders between clusters become less distinct and each individual cloud is more spread. Interestingly, we can associate nuclei with a distinct cell type even for those with as few as 100 genes detected (and without any requirement on the number of reads mapped to the given cell barcode), suggesting that the cell-type identity in the brain can be encoded by a small set of genes easily detected with shallow sequencing, as previously observed in other systems^9^. To visualize the expression of marker genes previously shown to be associated with specific sub-types of cells (*e.g.*, sub-types of GABAergic neurons in the hippocampus and cortex^3,20^), we visualized the average expression of the markers across each cluster and the distribution of the expression across cells in the tSNE space.

###### Testing for batch and technical effects

To validate that the resulting clusters are not driven by batch or other technical effects we checked the distribution of samples within each cluster and the distribution of the number of genes detected across clusters (as a measure of the nuclei quality). This was done both by visualizing the distribution along the tSNE space and by directly comparing the distributions of each of the above parameters between all clusters. The cells separated into distinct point clouds in tSNE space that were not driven by batch effects, each were an admixture of cells from all technical and biological replicates, with variable number of genes. Notably, there is a distinct difference in the number of genes between neuronal and glia nuclei in the brain, but cells cluster by cell type and not by the number of genes. However, when inspecting the tSNE maps more closely we do find that within some point clouds, there was a visible separation between nuclei according to the number of genes detected (but not by sample/batch), indicating that removing the first PC did not fully correct for these technical effects. In particular, cluster 11 in our human data (grey cluster in **Fig. 2a**) was associated mainly with one sample, and did not associate with specific cell type markers except for general neuronal markers, and expressed mitochondrial RNA, and thus it was discarded from further analysis.

###### Cluster annotation, filtering, differential expression, and pathway analysis

The major cell type clusters were identified by using a set of known cell type marker genes from the literature, as previously described^3,20^. In addition, we found for each cluster signatures of differentially expressed genes, which we used to further validate the identity of the cluster by matching these signatures with canonical cell type marker genes and by testing for enriched pathways. Differentially expressed signatures were calculated using student’s t-test between each pair of clusters, with FDR < 0.01 and requiring an average expression (number of unique UMIs) across all nuclei in the cluster to be bigger than log(1.1). We then consider a gene to be differentially expressed in a single cluster if it passes these thresholds in at least 60% of the pairwise comparisons. We consider a gene as a marker for a given cluster if it passes these thresholds in the pairwise comparison with all but one (valid) clusters, and in addition require a log-ratio>1.2 between the average expression across all nuclei in the given pair of clusters. The differential expression signatures and markers were tested for enriched pathways and gene sets using a hypergeometric test (FDR < 0.01). Pathways were taken from the MSigDB/GSEA resource (combining data from Hallmark pathways, REACTOME, KEGG, GO and BIOCARTA)^27^. We flagged as potential problematic clusters that are disregarded from downstream analysis, by two parameters: any cluster in which we could not find differentially expressed genes or enriched pathways; and clusters expressing overlapping markers of two different cells types that might be nuclei doublets. Several small clusters in the human and mouse brain analysis expressed overlapping markers of two different cells types, in most cases of neurons and oligodendrocyte cells, and were discarded (grey clusters, Figure **1f, 2a**).

---

**Figure S1.**
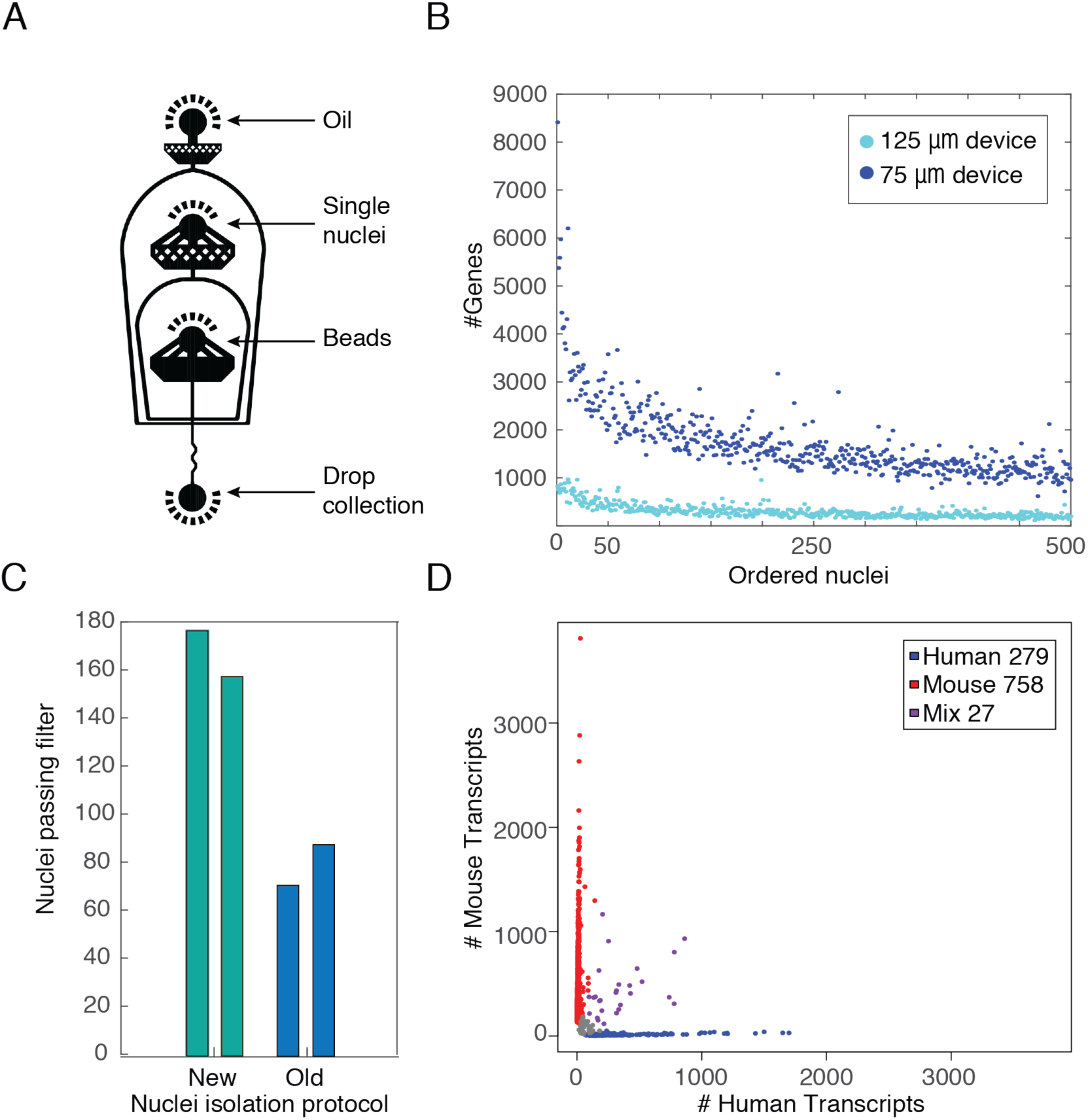
DroNc-Seq. (**a**) CAD schematic of DroNc-Seq microfluidics device. (b-d) DroNc-seq performance in 3T3 nuclei (channels are nominally 70 μm wide). (**b**) Number of detected genes. Scatter plot shows the number of detected genes (Y axis; defined as a gene with at least two different UMIs detected, **Methods**) across nuclei, ranked in decreasing order (X axis), using DroNc-Seq 75μm microfluidics device (blue, **Methods**) and Drop-Seq^5^ 125μm device (light blue). (**c**) Number of successfully profiled nuclei. Bar plot shows the number of nuclei passing library quality filter out of 1,288 (±114) nuclei per library (**Methods**, Y axis), using either sNuc-Seq^3^ (“old”) or DroNc-Seq (“new”) nuclei isolation protocol (**Methods)**. (**d**) Single nucleus specificity in DroNc-seq, estimated from mixtures of human 293 and mouse 3T3 nuclei. Scatter plot shows the number of UMIs associated with human (Y axis) or mouse (X axis) transcripts for each nucleus barcode (dot). Barcodes associated with a high number of both human and mouse transcripts (purple) reflect nuclei doublets. We find 2.5% (27/1,064) nuclei to be human-mouse, and thus estimate the expected doublet rate at our current loading and flow parameters to be 5%.

**Figure S2.**
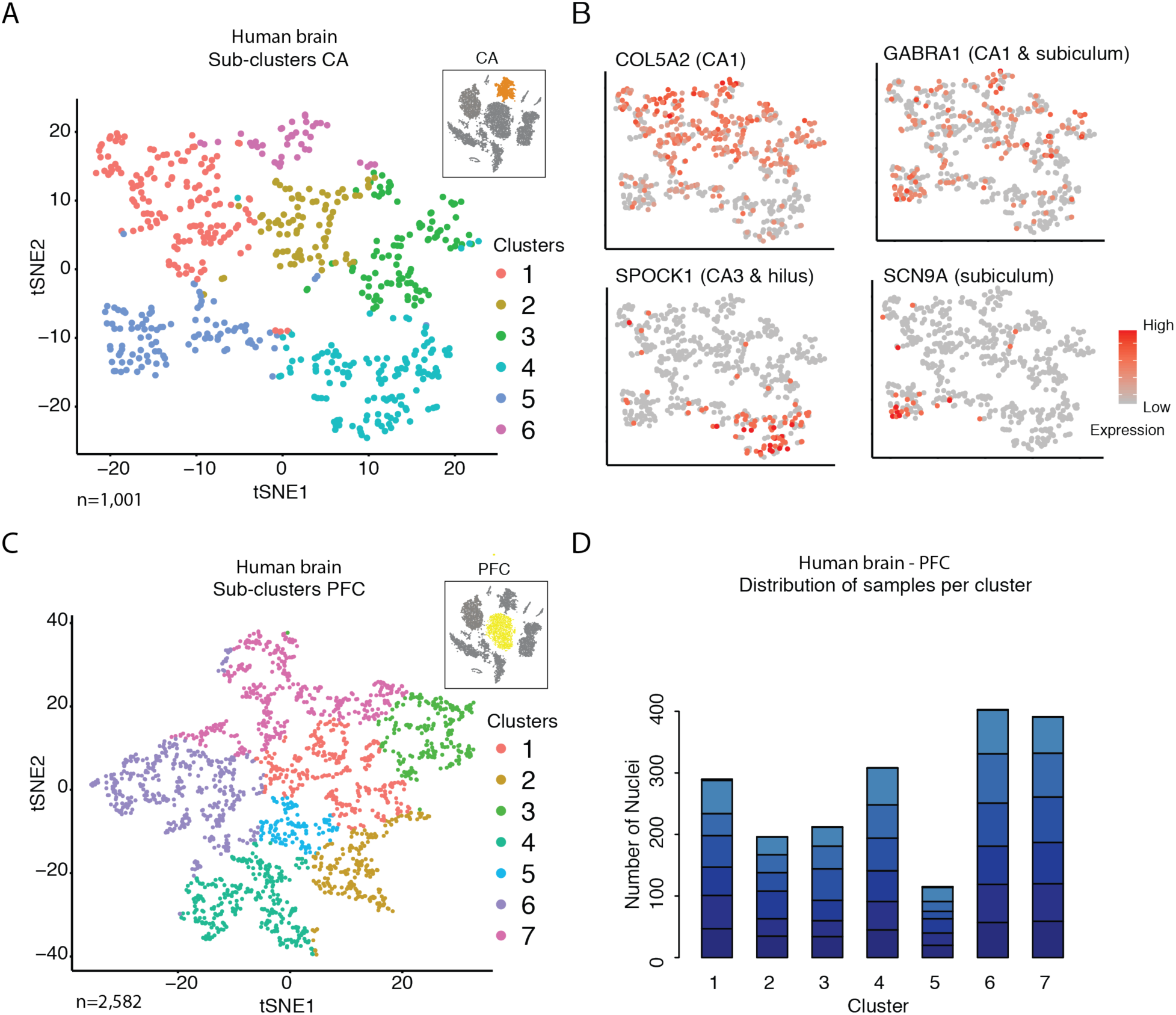
DroNc-Seq identifies sub-clusters for major cell types in the adult human brain. (**a,b**) Sub-clusters of the human CA pyramidal neurons cluster. tSNE embedding of DroNc-Seq profiles from the human CA pyramidal neurons cluster (cluster 2 in **Fig. 2a**; inset), color coded by sub-clusters (a) or by the expression of genes differentially expressed between the CA sub-clusters (b), with the known anatomical position of the gene according to the human Allen Brain Atlas^28^ noted in parentheses. Genes are: GABRA1 (highest in the CA1 and subiculum), SCN9A (subiculum), SPOCK1 (CA3 and hilus), and CL5A2 (CA1 based on the mouse Allen Brain Atlas^29^), showing that DroNc-Seq can distinguish between pyramidal neurons that are spatially separated along the anatomical sub-regions of the CA. (**c,d**) Sub-clusters of the human prefrontal cortex (PFC) pyramidal neurons cluster. (**c**) tSNE embedding of DroNc-Seq nuclei profiles from the human prefrontal cortex (PFC) pyramidal neurons cluster (cluster 1 in **Fig. 2a**, inset), color coded by sub-clusters. (**d**) Number of nuclei (Y axis) from each sample (color code) associated with each cluster (X axis), showing that each cluster is supported by multiple samples.

**Figure S3.**
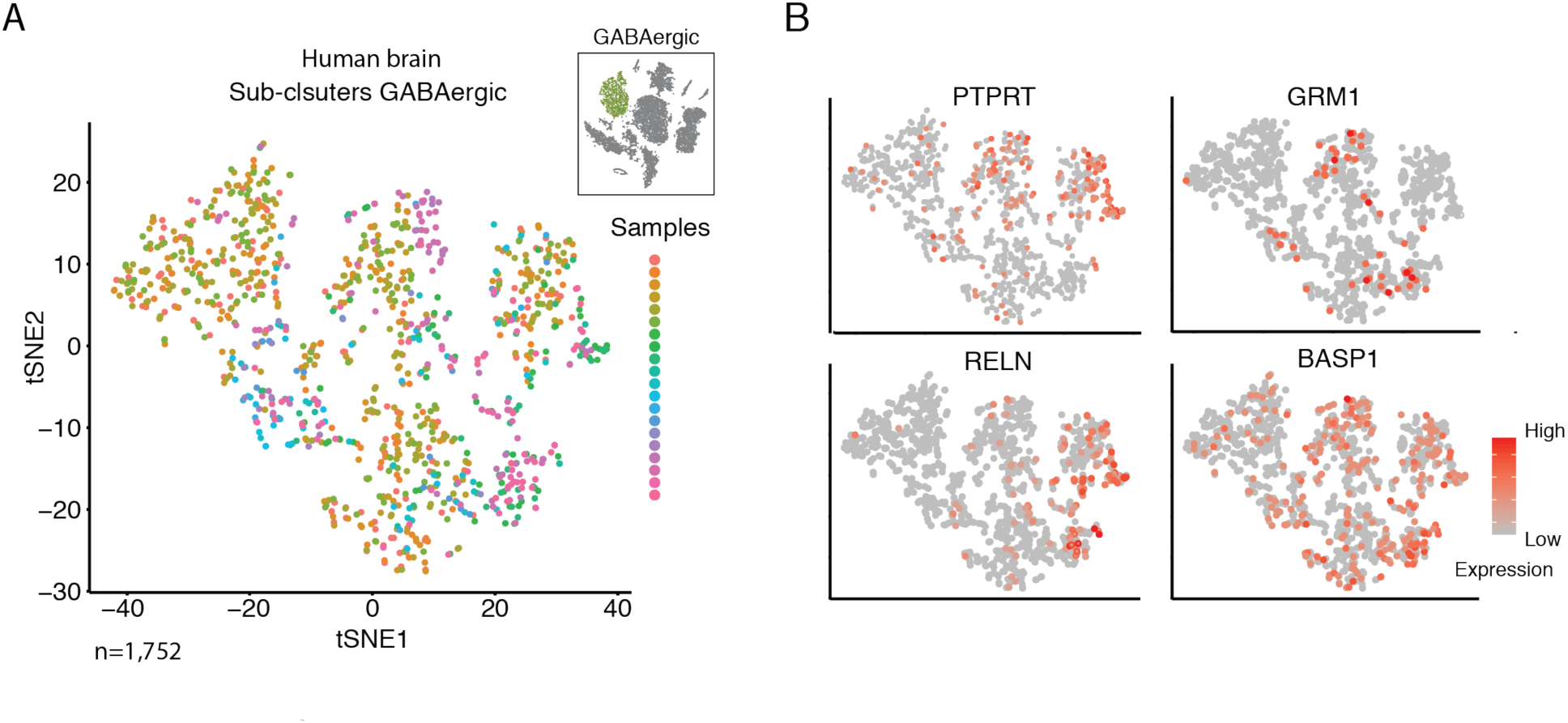
Sub-clusters of GABAergic neurons in the human brain. tSNE embedding (as in **Fig. 2g**) of DroNc-Seq nuclei profiles from the human GABAergic neuron cluster (cluster 3 in **Fig. 2a**, inset). (**a**) Color coded by the sample of origin, showing that each cluster is supported by multiple samples. (**b**) Colored by the expression level of known GABAergic marker genes or genes differentially expressed between the sub-clusters, showing unique combinatorial expression of patterns across clusters.

